# Scene complexity modulates degree of feedback activity during object recognition in natural scenes

**DOI:** 10.1101/293290

**Authors:** Iris I. A. Groen, Sara Jahfari, Noor Seijdel, Sennary Ghebreab, Victor A. F. Lamme, H. Steven Scholte

**Affiliations:** New York University, Department of Psychology, New York, NY, USA; University of Amsterdam, Department of Psychology, Section Brain and Cognition, Amsterdam, The Netherlands; University of Amsterdam, Department of Informatics, Intelligent Systems Lab, Amsterdam, The Netherlands

## Abstract

Object recognition is thought to be mediated by rapid feed-forward activation of object-selective cortex, with limited contribution of feedback. However, disruption of visual evoked activity beyond feed-forward processing stages has been demonstrated to affect object recognition performance. Here, we unite these findings by reporting that the detection of target objects in natural scenes is selectively characterized by enhanced feedback when these objects are embedded in high complexity scenes. Human participants performed an animal target detection task on scenes with low, medium or high complexity as determined by a biologically plausible computational model of low-level contrast statistics. Three converging lines of evidence indicate that feedback was enhanced during categorization of scenes with high, but not low or medium complexity. First, functional magnetic resonance imaging (fMRI) activity in early visual cortex (V1) was selectively enhanced for target objects in scenes with high complexity. Second, event-related potentials (ERPs) evoked by high complexity scenes were selectively enhanced from 220 ms after stimulus-onset. Third, behavioral performance deteriorated for highly complex scenes when participants were pressed for time, but not when they could process the scenes fully and thereby benefit from the enhanced feedback. Formal modeling of the reaction time distributions revealed that object information accumulated more slowly for high complexity scenes (resulting in more errors especially for fast decisions), and directly related to the build-up of the feedback activity that was observed exclusively for high complexity scenes. Together, these results suggest that while feed-forward activity may suffice for simple scenes, the brain employs recurrent processing more adaptively in naturalistic settings, using minimal feedback for sparse, coherent scenes and increasing feedback for complex, fragmented scenes.

**Author summary:** How much neural processing is required to detect objects of interest in natural scenes? The astonishing speed of object recognition suggests that fast feed-forward buildup of perceptual activity is sufficient. However, this view is contradicted by findings that show that disruption of slower neural feedback leads to decreased detection performance. Our study unites these discrepancies by identifying scene complexity as a critical driver of neural feedback. We show how feedback is enhanced for complex, cluttered scenes compared to simple, well-organized scenes. Moreover, for complex scenes, more feedback is associated with better performances. These findings relate the flexibility of neural processes to perceptual decision-making by demonstrating that the brain dynamically directs neural resources based on the complexity of real-world visual inputs.

## Introduction

Object recognition is often regarded as a task that is solved in the first wave of visual processing [1]. The human brain indeed recognizes objects at astonishing speed, with single neurons exhibiting object-selectivity 100 ms after stimulus onset [2], and global brain signals diverging within 100-200 ms [3,4]. Furthermore, hierarchical feed-forward models can emulate human performance [5,6], and neural representations in human and non-human primate brains match those in feed-forward neural [7–9].

However, the visual system does not conform to a strict feed-forward hierarchy: it contains long-range connections across hierarchical levels [10,11], many of which are feedback connections [12,13]. Feedback aids segmentation of figures from backgrounds [14–19] and perceptual completion [20] and is thought group scene elements together by implementing *visual routines* such as curve tracing and texture segmentation, which integrate individual line segments and other features encoded in low-level areas [21–23]. In accordance with this view, transcranial stimulation experiments have provided causal evidence indicating that both detection [24] and categorization [25] of target objects in natural scenes deteriorates when feedback is disrupted at time windows beyond feed-forward processing stages, that is beyond ~150 ms after stimulus onset.

How can we reconcile the speed of object recognition with an important role for feedback? Two lines of evidence suggest that feedback may be employed *adaptively* depending on the quality of the feed-forward sweep. First, computer simulations indicate that disrupting feedback has stronger effects for occluded or degraded target objects [26]. Second, backward masking, which interrupts feedback processing [27–29], is weaker for scenes with target objects that are “easily segregated” compared to “more demanding backgrounds” [30]. In this latter study, scene segmentation was assessed post-hoc, by asking observers to rate scenes on ‘ease of segregation’ of the target objects. However, scene perception research has shown that computational summary statistics may be diagnostic about scene complexity [31–33]. For example, the distribution of local contrast [34,35] can be estimated from simulated responses of early visual receptive fields using two summary parameters reflecting the contrast energy (CE) and spatial coherence (SC) in real-world scene [36,37]. These parameters describe a space in which sparse scenes separate from complex scenes with strong scene fragmentation (Figure 1*A)*. As these statistics appear to carry information about whether the scene contains objects that are clearly distinguishable from the background, we postulated that they could serve as useful predictors of the need for enhanced processing mediated by feedback.

**Figure 1.**
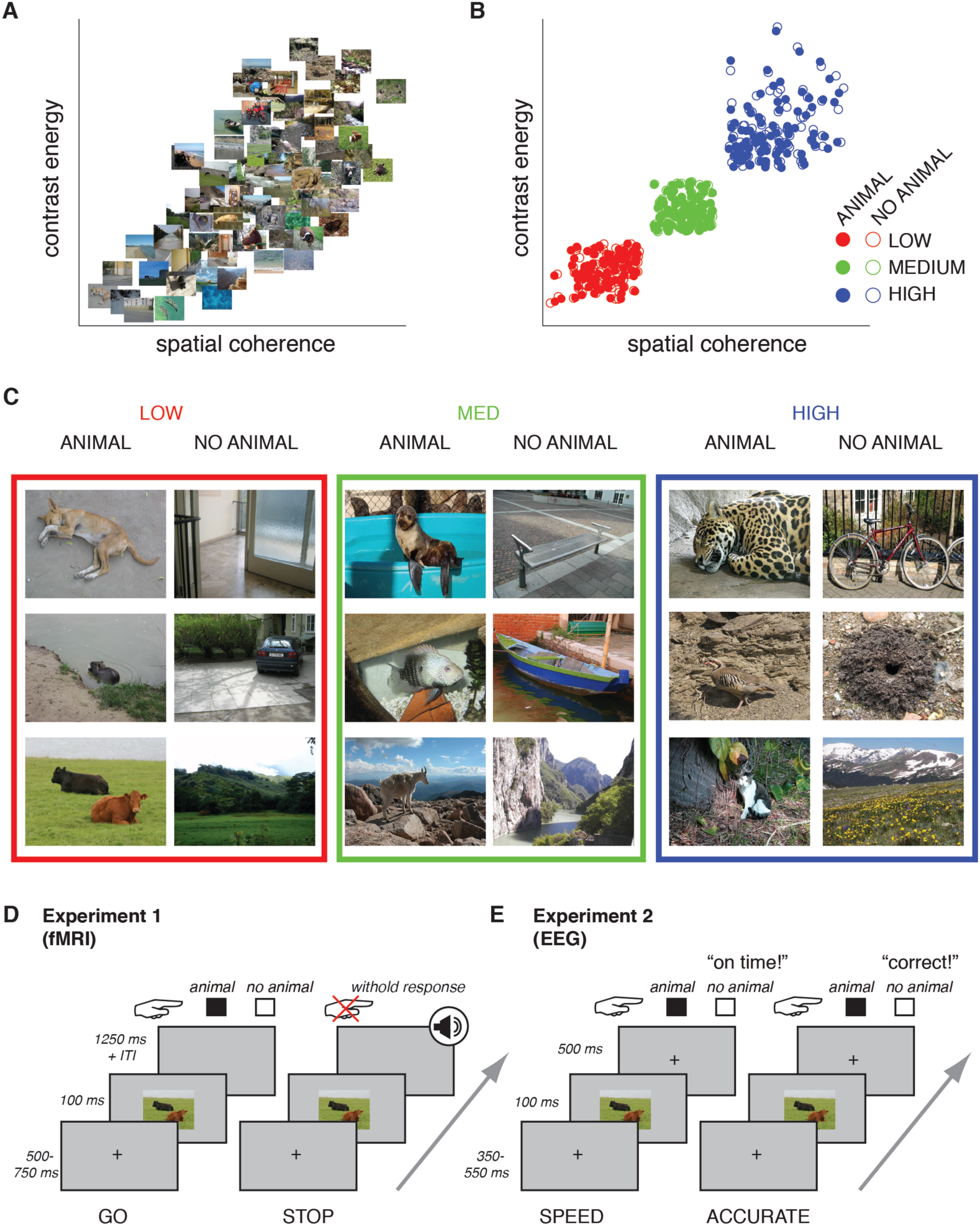
Stimuli and experimental paradigms. **A)** The diagnostic image statistics space described by two parameters derived from the distribution of local contrast, contrast energy (CE) and spatial coherence (SC). In this space, simple images containing one or a few easily segmentable objects are on the lower left, while complex images with a high degree of fragmentation are on the upper right. Thumbnails show 100 images (50 animal, 50 non-animal) randomly drawn from the larger image set from which the stimuli were selected. **B)** Image statistics of the stimuli: each point represents a scene sampled from the image space described in A). Scenes had either low (red), medium (green) or high (blue) CE and SC values. Within these conditions, CE and SC values were matched between scenes with target objects (animals, i.e. “A”; filled dots) and without target objects (non-animals, i.e. “NA”; open dots). **C)** Exemplars from each condition. **D)** Experimental design of the fMRI experiment. On GO trials, subjects indicated whether the scene contained a target object or not. On STOP trials, an auditory signal followed stimulus presentation after a variable inter-trial-interval (ITI), signaling that subjects had to withhold their response. Only GO trials were analyzed. **E)** Experimental design of the EEG experiment. Participants were instructed to respond as fast as possible on Speed trials and as accurate as possible on Accuracy trials and received feedback on each trial.

Here, we tested this hypothesis by measuring brain responses while participants performed a target object detection task in scenes that were systematically sampled to contain different summary statistics (see Figure 1*B-C*). Scene complexity was varied on a trial-by-trial basis, such that participants could not form an expectation of the scene complexity beforehand, allowing us to measure responses with unbiased feed-forward processing (i.e., without a difference in top-down task set or attentional state). First, we measured fMRI activity to target objects in scenes with different degrees of complexity (Experiment 1). Next, we measured EEG activity to the same scenes to examine the time-course of activity evoked by target objects (Experiment 2). Together, these experiments show that detection of target objects in high, but not low or medium complexity scenes was selectively associated with enhanced activity in early visual areas at late time-points in visual processing, indicating a feedback signal. Moreover, when participants emphasized response speed, decreases in behavioral performance were associated with a slowed evidence accumulation process (as formalized with a drift-diffusion model [38–40]) that was reflected in the late EEG activity (Experiment 2). Together, these results demonstrate a contribution of feedback to object recognition in complex scenes.

## Results

### Experiment 1 (fMRI)

#### Behavior

Behavioral performance for each condition is presented in Figure 2. Reaction times were found to increase gradually with scene complexity (F(2,44) = 4.96, p = 0.011, η^2^ = 0.18). Indeed, planned post-hoc comparisons showed the strongest increase for the HIGH complexity condition compared to the LOW (t(22) = 2.7, p(Sidák-corrected) = 0.035), and the MEDIUM condition (t(22) = 2.5, p(Sidák-corrected) = 0.057), with no significant difference between the LOW and MEDIUM conditions (t(22) = 0.7, p(Sidák-corrected) = 0.85; Figure 2A).

**Figure 2.**
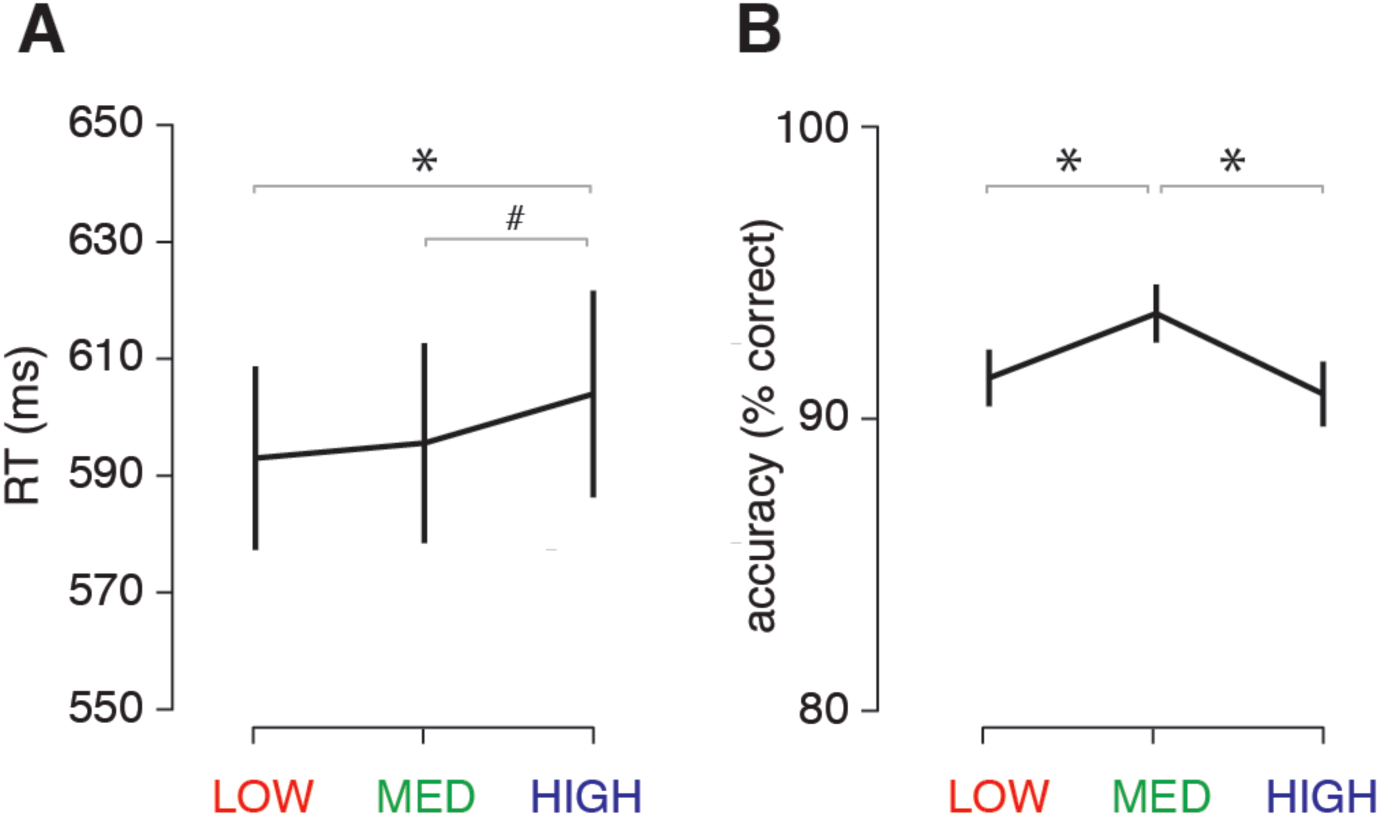
Behavioral results of the fMRI experiment. **A)** Average reaction times (RT) for the animal/non-animal categorization task per condition. **B)** Accuracy (percentage correct) per condition. Horizontal black lines indicate the statistical outcome of one-way repeated-measures ANOVAs; gray lines indicate the results of pairwise tests corrected for multiple comparisons using a Sidák correction. Error bars represent S.E.M. * = p < 0.05, # = p < 0.10.

Response accuracy was also modulated by scene complexity (F(2,44) = 8.37, p = 0.001, η^2^ = 0.28): post-hoc comparisons showed the best performance for the MEDIUM condition compared to the LOW (t(22) = 3.5, p(Sidák-corrected) = 0.007) and HIGH condition (t(22)= 3.2, p(Sidák-corrected) = 0.012), with no differences between the LOW and HIGH conditions (t(22)= 0.8, p(Sidák-corrected) = 0.77; Figure 2*B*), for which the most pronounced response time differences were found. These results indicate that the high scene complexity leads to significant slowing in decision-times. Although choice performance in the LOW and HIGH condition was worse than for the intermediate condition, responses were only slowed for HIGH complexity scenes, suggesting that the HIGH condition was experienced as most difficult.

#### fMRI results: whole brain analysis

Whole-brain comparisons of target-present (animal) vs. target-absent (non-animal) scenes revealed significant clusters in lateral and ventral high-level visual cortex, respectively (Figure 3). For the animal > non-animal contrast (Figure 3*A*), bilateral clusters overlying lateral occipital cortex were present in all conditions. Critically, in the HIGH condition, additional differential activity was present in low-level visual areas. Indeed, contrasting these statistical maps between conditions revealed a large cluster in several early visual areas (Figure 3*B*) and smaller clusters in inferior parietal regions. For the non-animal > animal contrast (Figure 3*C*), bilateral clusters in parahippocampal cortex were present in the MEDIUM condition: in the LOW and HIGH condition, only right-lateralized clusters survived whole-brain cluster-correction. Contrasting the difference between non-animal > animal scenes between conditions resulted in no significant clusters. Cluster coordinates for all contrasts are reported in Table 1.

**Table 1.**
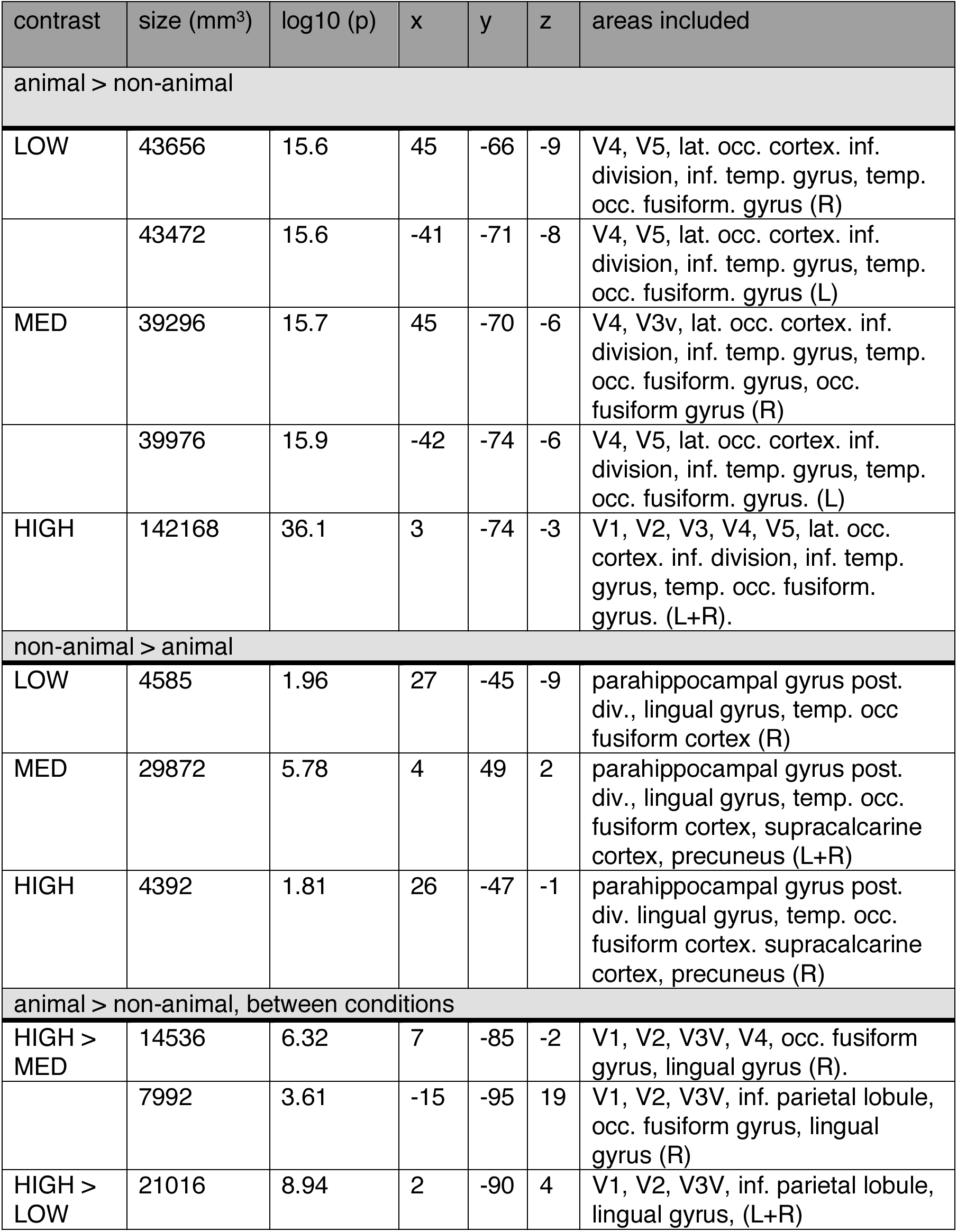
Whole brain fMRI analysis cluster coordinates for the significant contrasts. COG = center of gravity, p = p-value, lat. = lateral, occ. = occipital, temp. = temporal, inf. = inferior, post. = posterior, L = left, R = right. Coordinates are reported in MNI space. Areas included in the clusters were determined by determining overlap of local maxima within clusters with the probability maps of the Juelich histological Atlas and the Harvard-Oxford Cortical Structural Atlas implemented in FSL. Note that for the contrasts HIGH (animal - non-animal), MED (non-animal - animal) and [HIGH (animal-non-animal) - MED (animal-non-animal)], the clusters are bilateral, forming one cluster across hemispheres (see also Figure 3).

| contrast | size (mm<sup>3</sup>) | log10 (p) | x | y | z | areas included |
| --- | --- | --- | --- | --- | --- | --- |
| animal > non-animal |  |  |  |  |  |  |
| LOW | 43656 | 15.6 | 45 | -66 | -9 | V4, V5, lat. occ. cortex. inf. division, inf. temp. gyrus, temp. occ. fusiform. gyrus (R) |
|  | 43472 | 15.6 | -41 | -71 | -8 | V4, V5, lat. occ. cortex. inf. division, inf. temp. gyrus, temp. occ. fusiform. gyrus (L) |
| MED | 39296 | 15.7 | 45 | -70 | -6 | V4, V3v, lat. occ. cortex. inf. division, inf. temp. gyrus, temp. occ. fusiform. gyrus, occ. fusiform gyrus (R) |
|  | 39976 | 15.9 | -42 | -74 | -6 | V4, V5, lat. occ. cortex. inf. division, inf. temp. gyrus, temp. occ. fusiform. gyrus. (L) |
| HIGH | 142168 | 36.1 | 3 | -74 | -3 | V1, V2, V3, V4, V5, lat. occ. cortex. inf. division, inf. temp. gyrus, temp. occ. fusiform. gyrus. (L+R). |
| non-animal > animal |  |  |  |  |  |  |
| LOW | 4585 | 1.96 | 27 | -45 | -9 | parahippocampal gyrus post. div., lingual gyrus, temp. occ. fusiform cortex (R) |
| MED | 29872 | 5.78 | 4 | 49 | 2 | parahippocampal gyrus post. div., lingual gyrus, temp. occ. fusiform cortex, supracalcarine cortex, precuneus (L+R) |
| HIGH | 4392 | 1.81 | 26 | -47 | -1 | parahippocampal gyrus post. div. lingual gyrus, temp. occ. fusiform cortex. supracalcarine cortex, precuneus (R) |
| animal > non-animal, between conditions |  |  |  |  |  |  |
| HIGH > MED | 14536 | 6.32 | 7 | -85 | -2 | V1, V2, V3V, V4, occ. fusiform gyrus, lingual gyrus (R). |
|  | 7992 | 3.61 | -15 | -95 | 19 | V1, V2, V3V, inf. parietal lobule, occ. fusiform gyrus, lingual gyrus (R) |
| HIGH > LOW | 21016 | 8.94 | 2 | -90 | 4 | V1, V2, V3V, inf. parietal lobule, lingual gyrus, (L+R) |

**Figure 3.**
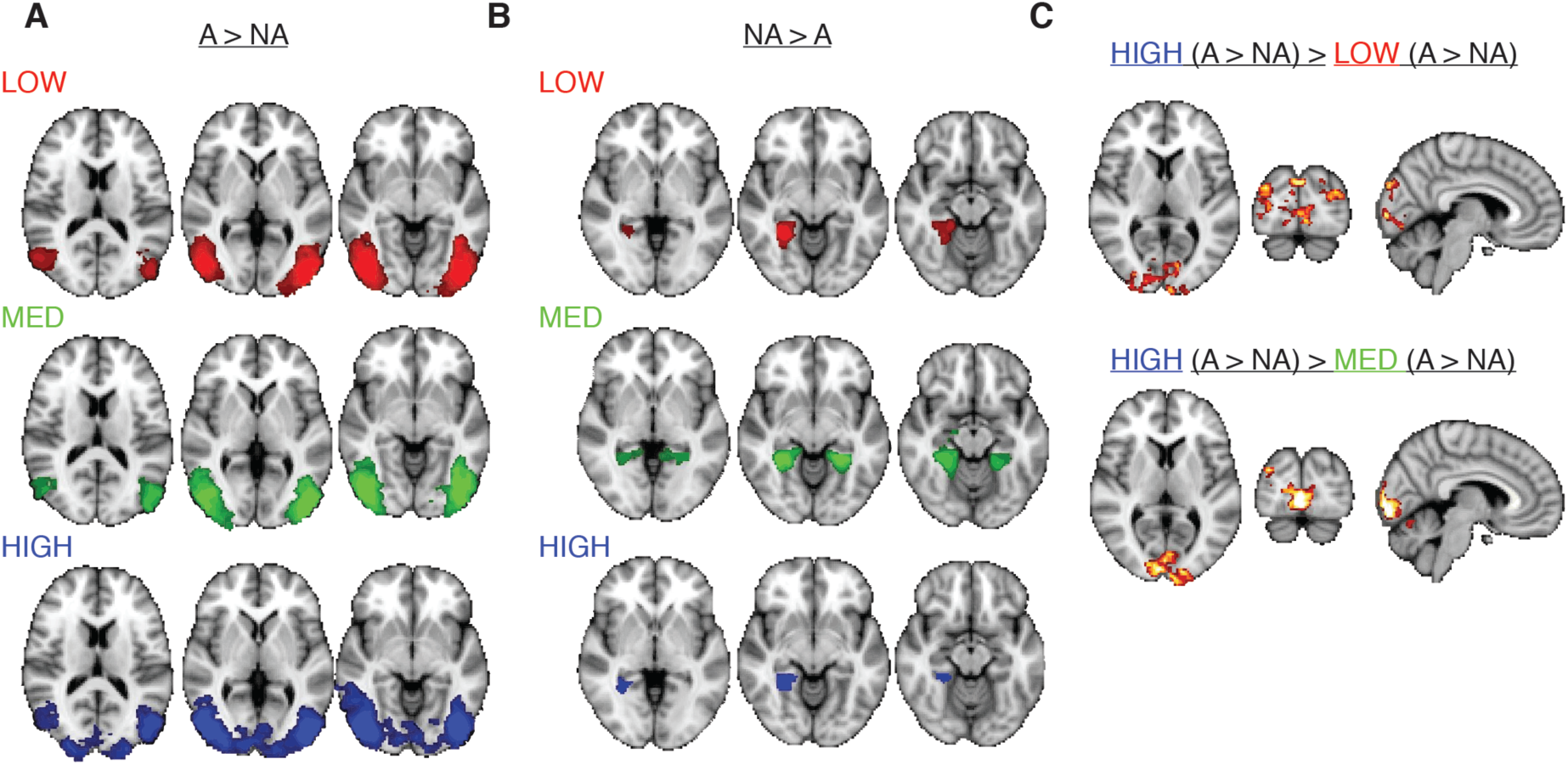
Whole brain fMRI results. **A)** Statistical parametric maps for the animal (A) > non-animal (NA) contrast for each condition. From left to right, MNI coordinates for each transversal slice are z = [10, 2, -6]. **B)** Statistical parametric maps for non-animal > animal contrast for each condition. From left to right, MNI coordinates are z = [-2, -8, -14]. **C)** Differences in the animal vs. non-animal activity between conditions. MNI-coordinates: [x = 6, y = -88, z = 6]. Maps were cluster-corrected and thresholded at z = 2.3. Color scales range from z = 2.5 to 5 (A and B) and z = 2.3 to 3.5 (C).

#### fMRI results: ROI analysis

Following the whole brain analysis, the difference in BOLD activity for animal vs. non-animal scenes in each condition was computed in four *a priori*, independently defined regions of interest and compared across conditions using repeated-measures ANOVAs. In line with the whole brain results, scene complexity was found to modulate activity in early visual cortex (V1: (F(2,44) = 4.9, p = 0.01, η^2^ = 0.18; Figure 3*A*. Planned post-hoc comparisons indicated that this effect was driven by increased activity for animal (vs non-animal) scenes in the HIGH compared with the MEDIUM condition (t(22) = 2.6, p(Sidák-corrected) = 0.05) and a trend towards larger differential activity in the HIGH versus the LOW condition (t(22) = 2.3, p(Sidák-corrected) = 0.09), with no difference between LOW and MEDIUM conditions (t(22) = 1.2, p(Sidák-corrected) = 0.54). Moreover, one-sample t-tests conducted for each of the three conditions indicated that the differential activity in V1 significantly deviated from zero only in the HIGH condition (HIGH: t(22) = 2.8, p = 0.03; LOW: t(22) = 0.19, p = 0.99; MEDIUM: t(22) = -1.28, p = 0.52, Sidák-corrected).

We also observed a main effect of scene complexity in place-selective PPA (F(2,44) = 3.6, p = 0.04, η^2^ = 0.14 (Figure 4*B*), which exhibited stronger responses for non-animal than animal scenes. Planned post-hoc comparisons indicated a trend towards less differential activity for the HIGH compared with the MEDIUM (t(22) = 2.4, p(Sidák-corrected) = 0.07), but not the LOW condition (t(22) = 1.9, p(Sidák-corrected) = 0.19), and no difference between the LOW and HIGH conditions (t t(22) = 0.7, p(Sidák-corrected) = 0.89). As expected, object-selective LOC and face-selective FFA both responded more positively to scenes with animals than to scenes without animals (Figure 4*C-D*), but these responses did not differ across conditions (FFA: F(2,44) = 1.7, p = 0.19, η^2^ = 0.07; LOC: F(2,42) = 1.9, p = 0.14, η^2^ = 0.08).

**Figure 4.**
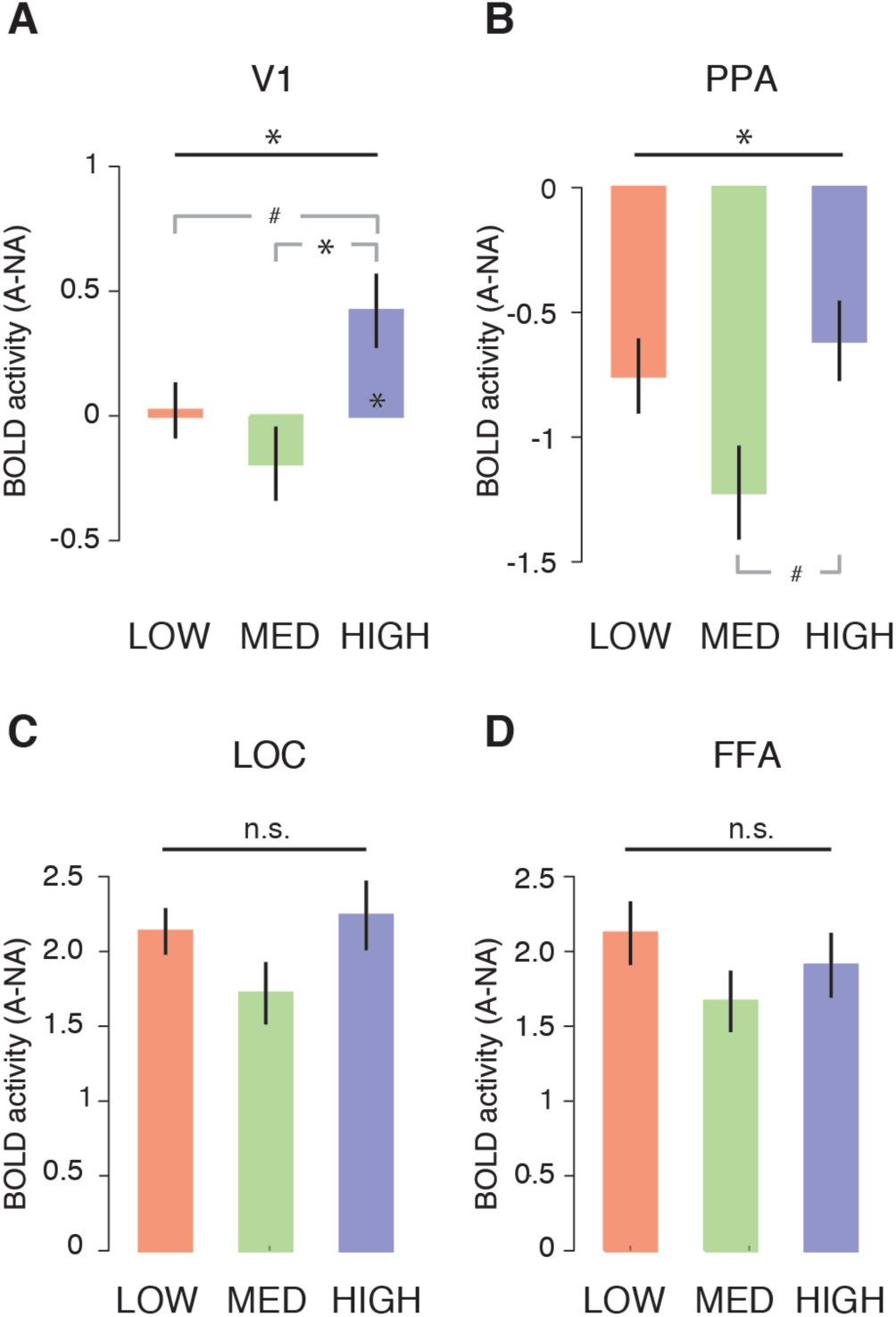
ROI analysis results for the fMRI experiment. Differential activity for animal vs. animal/non-animal scenes for each condition in **A)** V1, **B)** PPA, **C)** LOC and **D)** FFA. Horizontal black lines indicate the statistical outcome of repeated-measures ANOVAs; gray lines indicate the results of pairwise tests corrected for multiple comparisons using a Sidák correction. * = p < 0.05, # = p < 0.10. Error bars represent S.E.M.

In sum, these indicate that in all three conditions, target presence was reflected in known object- and face-selective regions, which were not significantly modulated by scene complexity. In contrast, scene complexity modulated activity levels in V1 and scene-selective PPA. In PPA, differential activity related to target-presence was least pronounced for the high complexity scenes compared to the medium complexity scenes, while V1 activity was only modulated by target presence for high complexity scenes. Taken together, these results indicate that scenes with high complexity induced the least difference in PPA while giving rise to additional differential activity in early visual areas.

#### Summary Experiment 1

The fMRI results suggest that information encoded in V1 may be selectively recruited during target object detection for the most difficult, high complexity scenes. Since the scenes were carefully matched in CE and SC *within* complexity levels, it is unlikely that this activity was driven by differences in low-level properties between target and non-target scenes evoked by feed-forward processing. Therefore, we hypothesized that the differential V1 activity results from increased feedback elicited by the need for increased processing of visual information related to the target object in high complexity scenes.

However, two limitations preclude a strong confirmation of this hypothesis. First, due to the poor temporal resolution of the fMRI signal, we cannot exclude the possibility that early visual activity differences were present as a result of other differences between the target and non-target scenes than those captured by CE and SC. Second, the categorization task was embedded in a stop-signal paradigm (see Materials and Methods). Although all results reported above were based on GO trials only (i.e. omitting all trials in which a stop signal was presented), the observed effects could be specific to, or amplified by, an expectation of a stop instruction, and might not generalize to a task in which participants are never asked to withhold their response.

To overcome these two limitations, we conducted a second experiment, in which we recorded EEG while participants performed an animal/non-animal categorization task on the same set of scenes without any stop-signal manipulation. First, to test our hypothesis that the activity differences in early visual areas were feedback-related, we computed event-related potentials (ERPs) for target and non-target scenes separately in each condition, to infer when these responses started to differ on electrodes overlying visual cortex. Again, since scenes were matched in image statistics, we predicted that there would be no differences between animal and non-animal scenes before ~150 ms after stimulus onset, beyond which feedback starts to affect ERP activity [23,28,41]. Second, because categorization of the high complexity scenes was characterized by slowed decision times with no substantial improvements in accuracy, the behavioral task in the EEG experiment was divided into blocks with a ‘speed’ or ‘accuracy’ emphasis. We hypothesized that if target detection in high complexity scenes was indeed associated with increased feedback, performance in the HIGH condition should be most affected for speeded trials, in which extensive visual processing is limited by time constraints.

### Experiment 2 (EEG)

#### Behavior and HDDM parameters

Figure 5 shows an overview of the behavioral and modeling results in Experiment 2. Analysis of the reaction times showed a significant main effect for instruction (‘speed’, ‘accurate’; F(1,25) = 87.6, p < 0.001, η^2par^ = 0.78), and scene complexity (‘LOW’, ‘MEDIUM’, ‘HIGH’; F(2,50) = 29.1, p < 0.001, η^2par^ = 0.54). As expected, participants responded faster with speed instructions, and response times were again longest for the high complexity scenes. Planned post-hoc comparisons further showed that responses were significantly slower for high compared with medium complexity scenes under both instructions (speed: t(25) = 5.0, p < 0.001; accurate: t(25) = 5.1, p = < 0.001; all Sidák-corrected; Figure 5*A*). Compared to low complexity scenes, responses were significantly slower for accurate task instructions (t(25) = 4.9, p > 0.001), while showing a similar trend for the speed instruction (t(25) = 2.7, p = 0.065; all Sidák-corrected). Response times did not differ between low and medium conditions (speed: t(25) = 2.4, p = 0.13; accurate: t(25) = 0.72, p = 0.98; all Sidák-corrected).

**Figure 5.**
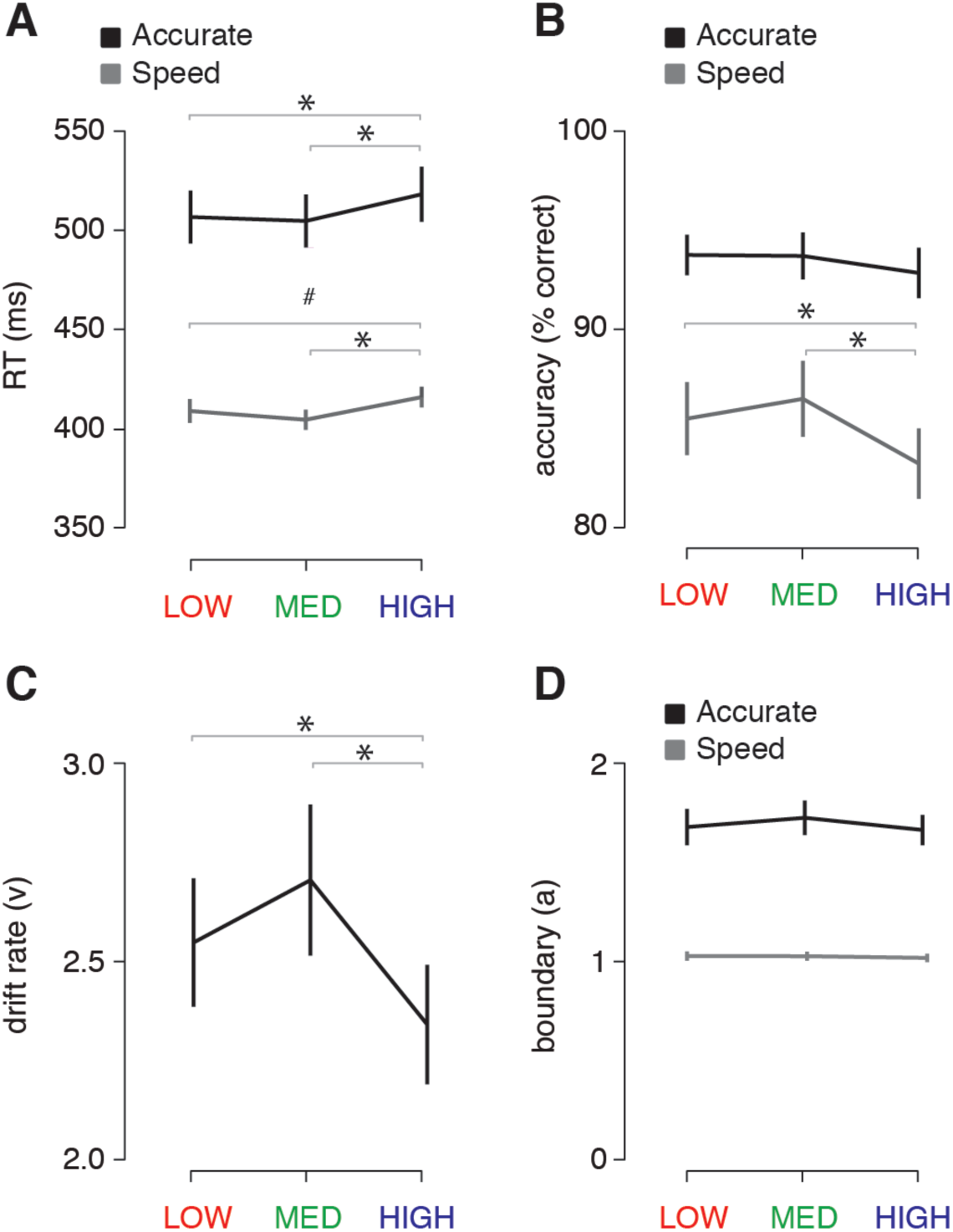
Behavioral and modeling results of the EEG experiment. **A)** Average reaction times (RT) per condition and task instruction. **B)** Accuracy (percentage correct) per condition and task instruction. **C)** Drift rates per condition, indicating slower evidence accumulation for high complexity scenes. **D)** Estimates of the amount of evidence required per condition and task instruction. Gray lines indicate the results of pairwise post-hoc tests between conditions corrected for multiple comparisons using a Sidák correction. Error bars represent S.E.M. * = p < 0.05.

Next, the inspections of accuracy scores showed a significant interaction between scene complexity and task-instruction (F(2,50) = 5.6, p = 0.002, η^2par^ = 0.18). That is, detection accuracy was selectively impaired for highly complex scenes only when participants were motivated to respond as fast as possible (HIGH vs. LOW, t(25) = 3.2, p = 0.024; HIGH vs. MEDIUM, t(25) = 4.4, p = 0.001; LOW vs. MEDIUM, t(25) = 1.98, p = 0.38, all Sidák-corrected). Critically, this effect was not found for trials in which participants were asked to be as accurate as possible (HIGH vs. LOW; t(25) = 1.5, p = 0.60; HIGH vs. MEDIUM, t(25) = 1.8, p = 0.42; LOW vs. MEDIUM: t(25) = 0.1, p = 0.99, all Sidák-corrected; Figure 5*B*).

In a final step, we estimated HDDM parameters to better understand the prolonged RT’s observed when participants were presented with highly complex scenes, as well as the selective decrease in detection accuracy when participants were pressed for time. Consistent with the observations above, the speed of information accumulation (drift rate, v) was modulated across the three scene complexity conditions (F(2,50)=12.5, p<0.001, η^2par^ = 0.33), with the slowest rate of information accumulation for highly complex scenes (HIGH vs. LOW; t(25) = 2.97, p = 0.02; HIGH vs. MEDIUM, t(25) = 5.04, p < 0.001; LOW vs. MEDIUM: t(25) = 2.05, p = 0.15, all Sidák-corrected; Figure 5*C*). In contrast, the amount of evidence that is required to make a choice was only adapted as a function of task instructions (F(1,25) = 93.5, p<0.001, η^2par^ = 0.79), and was not modulated by scene complexity (F(2,50) = 1.3, p = 0.22, η^2par^ = 0.05), or an interaction (F(2,50) = 2.5, p = 0.10, η^2par^ = 0.09; Figure 5*D*).

Together, these results replicate the findings from Experiment 1, and furthermore show: 1) prolonged reaction times for highly complex scenes, with 2) more errors when the decision is speeded, because of 3) a slower rate of information processing.

#### ERP results

To investigate the time-course of object categorization in visual cortex, evoked responses to the target and non-target scenes were pooled across a set of occipital and peri-occipital electrodes (see Materials and Methods). The results in Figure 6 clearly show that while in the LOW and MEDIUM conditions, target and non-target ERPs diverged at only a few time-points across the entire epoch, the waveforms in the HIGH condition were robustly enhanced for animal scenes at occipital electrodes starting from ~220 ms after stimulus onset. These effects were observed for both the speed and accuracy trials.

**Figure 6.**
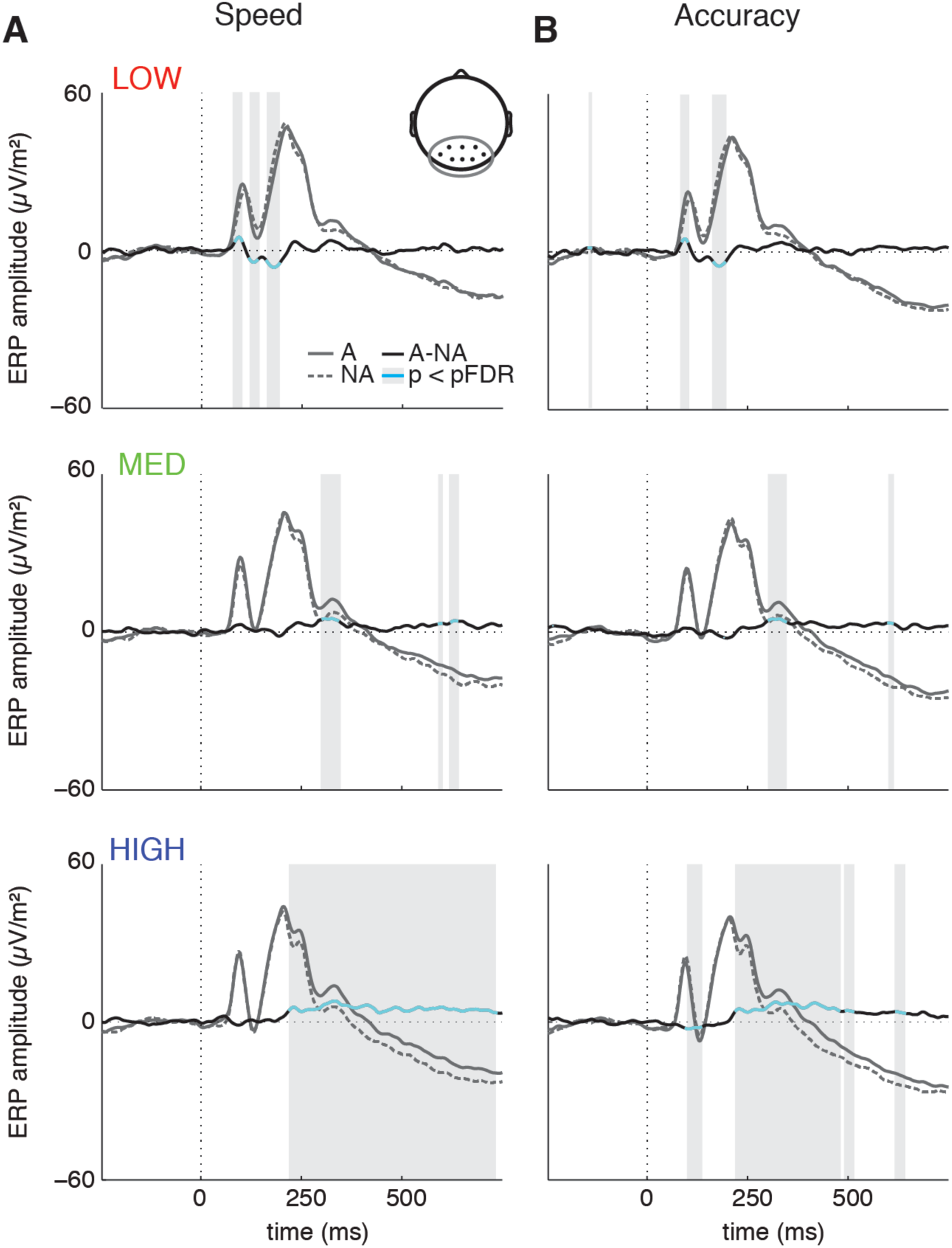
ERP results. Average ERP amplitude for animal (solid gray lines) and non-animal scenes (dashed gray lines) and the corresponding difference wave (solid black lines) per condition for an occipital-peri-occipital cluster of EEG channels (Oz, POz, O1, O2, PO3, PO4, PO7, PO8) for the **A)** speed and **B)** accuracy trials. Significant difference waves (FDR-corrected across tasks, conditions and time-points) are indicated with gray boxes and cyan lines.

Statistical comparisons (FDR-corrected across all time-points, task-instructions and conditions) of each of the differences waves against baseline indicated that for speeded trials, the difference wave in the HIGH condition was insignificant before 220 ms (Figure 7*A*, blue solid line); in the accuracy trials, there was a brief interval of significant deflection before 200 ms (Figure 7*B*), but this reflected a reversed difference wave compared to the other two conditions. Beyond 220 ms, however, this pattern reversed: direct comparison of the difference wave across conditions indicated that the difference wave for the HIGH condition after 220 ms was significantly enhanced compared to the LOW (blue asterisks; top) and MEDIUM (plusses; top) conditions.

**Figure 7.**
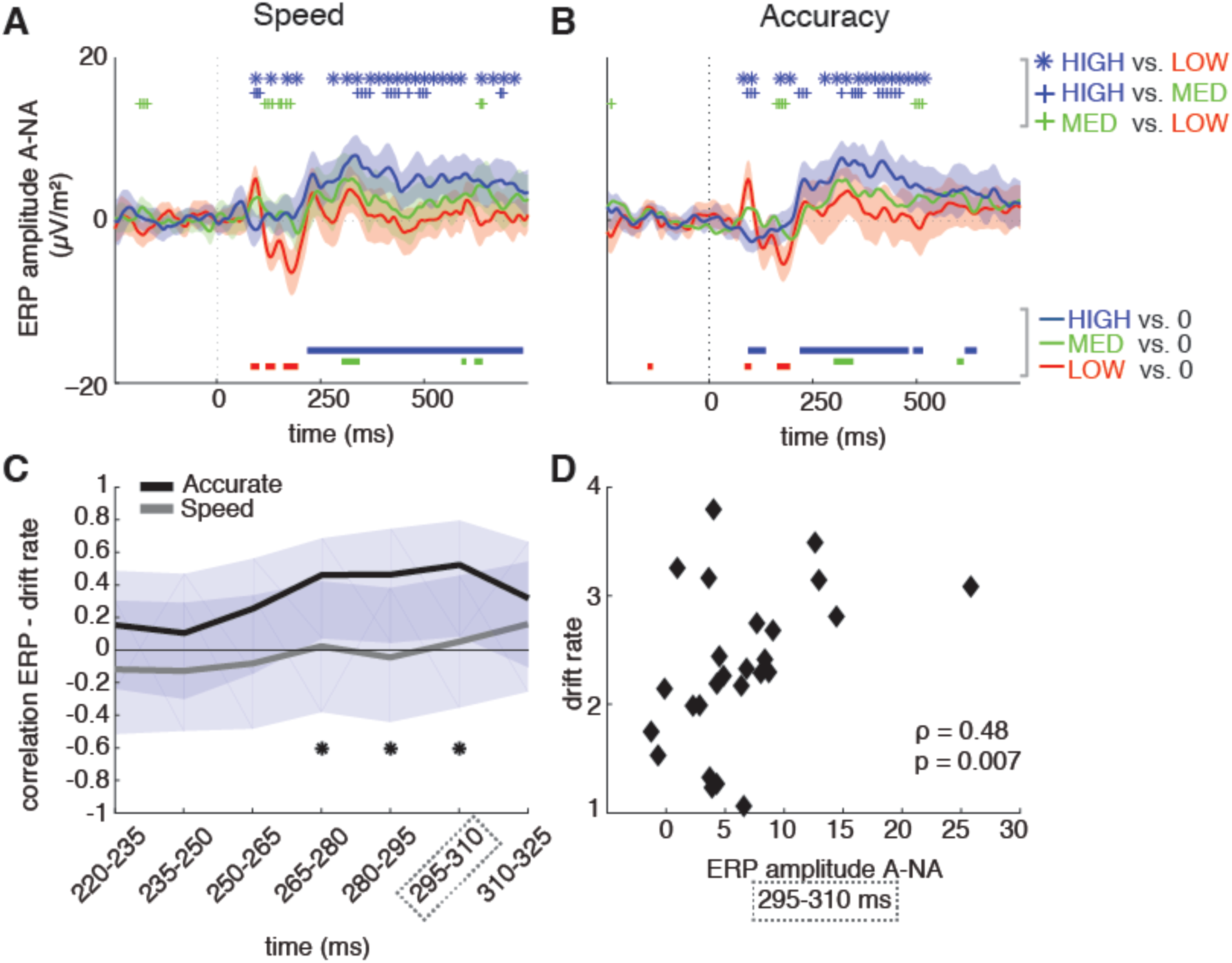
Difference wave comparison and correlation with behavior. Difference wave comparisons (FDR-corrected across time-points, task-instructions, conditions and tests) for each condition overlaid and statistically compared for the speed **A)** and accuracy **B)** trials. Thick lines at the bottom of the graphs indicate significant difference wave deflections from zero, and shadings indicate 95% confidence intervals derived from one sample t-tests. Symbol markers indicate significant differences in difference wave amplitude between conditions, blue asterisk: HIGH vs. LOW; blue plus: HIGH vs. MEDIUM; green plus: MEDIUM vs. LOW. **C)** Correlations of difference wave amplitude in the HIGH condition with drift rate (Spearman’s rho, FDR-corrected across time-points) for 15 ms intervals between the first significant deflection (220 ms) until peak (325 ms). Asterisks indicate significant correlations; shadings indicate 95% confidence intervals derived from a percentile bootstrap (10.000 iterations). **D)** Drift rates for each participant in the HIGH condition compared with their difference wave amplitude in the HIGH condition at the interval of highest correlation (295-310 ms) (see dashed box in panel C).

The absence of an early difference wave for the high complexity scenes aligns with the behavioral observations, where categorization was mostly affected under time pressure and could not profit from the substantial differentiations later in time. These results further show that it is unlikely that the differential fMRI activity in early visual areas in the HIGH condition is driven by uncontrolled image properties. Instead, we observed a strongly enhanced ERP difference beyond 200 ms in this condition, suggesting an electrophysiological correlate of the early visual area difference observed in the fMRI experiment. Conversely, these results suggest that EEG activity before 200 ms contributed little, if anything, to the V1 differences observed in the fMRI data.

#### Relating ERPs to behavior

The EEG results demonstrated that beyond 200 ms, the feedback signal is similarly enhanced for both speed and accurate instructions, whereas detection accuracies for the highly complex scenes were only impaired during speeded decision trials (for which there is no time to wait for the slow incoming information). Therefore, we hypothesized that the observed feedback during speeded trials, while present, did not benefit information processing. To test this hypothesis, we correlated difference wave amplitude in the HIGH condition (from the time at which the difference waves deflected significantly from zero) to the estimated speed of information processing (drift rate) for the high complex scenes. Critically, this evaluation showed a positive relationship between difference wave magnitude and the speed of information processing for accurate instruction trials, but not speeded trials (Figure 7C. This correlation was most pronounced just prior to the peak difference wave amplitude at 325 ms, i.e. around 300 ms after stimulus onset (Figure 7D), suggesting a ‘ramping up’ of activity commensurate with the amount of evidence accumulated (drift rate) for the animal/non-animal decision.

#### Summary Experiment 2

The ERP signal related to target presence in the HIGH, but not the LOW and MEDIUM conditions, was strongly enhanced after 200 ms, suggesting that it cannot arise from a feed-forward signal alone, but is modulated by feedback. This late enhancement was present for all trials (speeded or accurate) but only correlated with the rate of evidence accumulation for accurate trials, suggesting that behavioral performance benefits when participants are given sufficient time to process this signal.

## Discussion

We investigated the influence of scene complexity on successful detection of target objects in natural scenes using fMRI, EEG and behavior. Behaviorally, we observed prolonged decision times when participants viewed high compared with low complexity scenes. Using fMRI, we found this slowing to be accompanied by a selective increase in activity for target objects in early visual regions extending as far back as V1, while target activity in higher-level object-selective cortex was not affected by scene complexity. EEG recordings suggested that the increased activity across early visual cortex is likely driven by feedback arising after ~200 ms. While increased feedback for target objects in highly complex scenes was observed for both fast and accurate task instructions, behavioral performance only profited when the instructions emphasized accuracy, and so encouraged participants to wait for evidence accumulated via feedback.

We interpret these findings as showing a selective need for increased detailed analysis for the scenes of high complexity. For simple scenes, the initial, coarse representation provided by the feed-forward sweep at the beginning of the trial is enough for high-level areas to obtain information about the presence or absence of a salient object. For highly complex scenes, however, the feed-forward sweep is not sufficiently informative: it merely signals that there is a lot of ‘stuff’ in the scene, and detecting the target object requires additional processing. This interpretation fits with the global-to-local processing framework whereby detailed scene analysis takes place via reentrant processing [23,42–44]. In this framework the feed-forward sweep provides visual cortex with a *base representation.* If the visual task can be solved based on this representation alone, no further operations are necessary and action selection can be initiated. If it is not sufficiently informative, *elemental operations* are applied such as contour grouping, curve tracing and texture segmentation, which integrate individual line segments and other features encoded in low-level areas via incremental grouping [21,22].

While this theoretical framework is phrased in neurophysiological terms, a psychological correlate of this feedback-driven process could be termed ‘attentive processing’ while the base representation created in the feed-forward stage could be considered pre-attentive, potentially reflecting different modes of visual processing that are components of psychological theories such as Feature Integration Theory [45] that distinguish pre-attentive parallel processing across the entire scene (“feed-forward”) from conjunctive processing via an attentional spotlight (‘feedback/recurrent”); see Roelfsema, (2006) for further discussion about similarities and differences between these frameworks). Our observation of increased fMRI and EEG responses for targets is clearly consistent with the numerous observations that attention is reflected in response enhancements in visual cortex [46]. Importantly, however, our results suggest that this response enhancement is *selectively applied* on a trial-by-trial basis to scenes which have high complexity, as determined by bottom-up, biologically plausible image statistics.

### A role for image statistics in signaling the need for feedback

Our results suggest that the complexity of the feed-forward representation can be estimated using global image statistics. These statistics are suitable computational substrates for such a representation because they can be computed directly from local contrast responses in LGN [34]. These parameters strongly affect the amplitude of evoked EEG activity early in time in visual processing [34,37,47] and could therefore serve as ‘markers’ to determine whether further operations are necessary. This idea is reminiscent of previous proposals suggesting that scenes are summarized as coarse ‘blobs’ that precede detailed analysis at smaller spatial scales depending on their diagnostic value [48,49], or via low spatial frequencies that are used to direct top-down facilitation [50]. Flexible employment of feedback depending on the diagnosticity of the input is also consistent with results showing that feedback is necessary to model categorization of degraded scenes [26] and with results reported by [30], who found that masking effects were stronger for scenes that were ‘less easily segmented’, (as assessed post-hoc). Here, rather than filtering, degrading or otherwise manipulating the scenes, we used biologically plausible scene statistics to sample real-world variation in scene complexity, providing an a priori, quantitative approach to estimate this property.

### The impact of feedback on behavior

Our behavioral observations concur with selective feedback requirements in two important ways. First, in the fMRI experiment we observed that scene complexity influenced response selection (which requires active categorization), but not response suppression [51–54]. Specifically, we observed that the decision time was prolonged for complex scenes, whereas accuracy was similar to simple scenes. The prolonged decision times suggest a slower rate of information accumulation information for complex scenes. Indeed, the increased V1 activity further implied that for complex scenes, fast feed-forward representations alone might not be sufficient for optimal scene categorization. In the EEG experiment we examined this hypothesis through the manipulation of time restrictions. Behaviorally, we showed that while RTs were always increased for highly complex scenes, accuracy rates only declined when the instructions emphasized a speeded response. The well-known drift diffusion model [38,39,55] was then used to show how the rate of information processing was slowed for highly complex scenes [56–58], and indeed had a negative effect on target detection performance when participants were encouraged to make speeded decisions (as thus not wait for the slower incoming information). We observed that the speed of information processing (drift rate) was only related to the ERP difference wave when participants emphasized accuracy above speed (and thus were able to process the feedback signal). The ramp-like characteristic of this relation is consistent with previous ERP findings of a late ‘discriminating component’ around 300 ms reflecting the amount of sensory evidence for perceptual decisions on objects in phase noise [57,59]. Taken together, these observations underline the importance of the slow feedback for processing complex scenes to optimize behavioral performance.

### Where does the feedback come from?

A question that remains unresolved in this study is which, if any, brain areas might be involved in employing neural feedback based on scene complexity. Consistent with previous studies [60,61], we found a strong distinction between animal- and non-animal images in object-selective LOC and face-selective FFA. However, this differential activity was not strongly modulated by whether the object is embedded in a complex scene or a simple scene. We did observe, however, an effect of scene complexity in PPA. While the scene-selectivity of PPA is commonly attributed to coding of 3D spatial layout [62–64] it is also sensitive to object information [65–67], as well as low-level features such as spatial frequency, contrast, rectilinearity and texture [67–72]. suggesting that PPA may be a suitable region to demonstrate an influence of the broader scene context on object-related activity. Importantly, PPA is biased towards the visual periphery [73,74], containing relatively large, peripheral receptive fields [75,76] making it suited for computation of larger-scale summary statistics of the input [77]. Interestingly, a recent study found that PPA represented scene complexity reflected in various image-computable computational measures such as image compression and self-similarity [78]. Thus, one explanation of our results could be that the enhanced feedback was initiated by a feed-forward, summary statistics representation in PPA indicating a lack of sufficient differential object information for high complexity scenes.

This interpretation is highly speculative, however; the effects in PPA were relatively weak, and we do not know whether these PPA effects arise early or late in visual processing. The feedback signal could also originate, for example, in LOC or other high-level regions [20,79], but not be detectable with fMRI due to the temporal smoothing of the signal, causing these regions to only represent the ‘end result’ of the categorization (i.e., successful detection of an animal). Another possible scenario is that the enhanced activity does not originate from higher-level category-selective regions but from local recurrent circuits [21] within earlier visual areas, for example based on computations of local statistical dependencies in areas V2-V4 [80,81].

In addition, scene complexity is obviously not the only factor that influences the employment of feedback. Top-down attentional manipulations affect ERP responses to natural scenes in feedback time windows [47], and feedback is involved in the application of attentional templates [82,83], which may differ between different tasks [84] or the level of detail necessary to solve the task [49] and therefore in the amount of feedback required. While we varied scene complexity on a trial-by-trial basis, making it difficult for participants to use a top-down strategy to ‘predict’ how much attention they would need to direct to solve that trial, they still needed to apply a search target in order to solve the task. This search target was the same throughout the entire experiment (animal), but our whole-brain fMRI analysis (Figure 3) did indicate some target-related activity differences in posterior parietal regions for high complexity scenes, a region that has been associated both with holding representing attentional templates and representing outcomes of attentional selection [82]. A deeper understanding of the interaction between attentional template, bottom-up scene complexity and target-related feedback activity and their respective representation in cortical regions requires future experimental study. So far, however, our results suggest that although object recognition based on feed-forward information [5] is possible for certain scene types and tasks, in real-world perception there is a bidirectional interplay between bottom-up processing and top-down requirements [85], affecting dynamic feedback employment.

## Materials and Methods

### fMRI experiment

#### Subjects

Twenty-five participants (7 males, age 19-26 years; mean = 21.6, SD = 1.7) took part in the fMRI experiment. All participants had normal or corrected-to-normal vision, provided written informed consent and were financially compensated. The experiment was approved by the ethics committee of the University of Amsterdam. One participant had a median reaction time of 2 standard deviations above average as well as 7.5% non-responses and was therefore excluded from further data analysis. Another participant was excluded because of excessive head movement (absolute displacement of twice the voxel size caused by multiple movements across a single experimental run).

#### Stimuli

Scenes were selected from a larger set of 4800 scenes used in a previous EEG study [86]. The larger dataset contained scenes from several online databases, including the INRIA holiday database [87], the GRAZ dataset [88], ImageNet [89], and the McGill Calibrated Color Image Database [90]. For each scene, one CE and one SC value was computed by simulating the output of contrast-sensitive receptive fields and integrating these responses across the scene by averaging (CE) and divisive normalization (SC). In natural scenes, CE and SC correlate strongly with parameters of a Weibull function fitted to the distribution of contrast values [34,35], which is informative about the degree of scene fragmentation [91]. CE is a biologically realistic approximation of the distribution width (the scale parameter of the function), informing about the overall presence of edges in an image, whereas SC is an approximation of its shape (the degree to which the function describes a power law or a Gaussian distribution), capturing higher-order correlations between edges. As a result, images with low CE/SC values have strong ‘Inherent’ figure-ground segmentation, often containing a large (central) object surrounded by a homogenous background (and therefore containing few edges that are spatially correlated because they belong to the same object), whereas images with high CE/SC values are more complex, and typically textured or cluttered (with many edges distributed in a Gaussian manner that are uncorrelated); while they can contain large objects, they often have a background consisting of uncorrelated edge ‘noise’ (see also De Cesarei et al., (2017), footnote 4). The model is described in more detail in [37,47].

Here, we used these image statistics to selectively sample scenes with various levels of complexity. We created three conditions: LOW, MEDIUM and HIGH (Figure 1*B*), whereby each condition was defined by its CE/SC values. For the LOW and HIGH condition, we selected the lower and upper 25% percentiles of the distribution of CE/SC values in the full stimulus set, respectively; for the MEDIUM condition, the middle 35% percentile. Each condition consisted of 160 images, half of which contained an animal. Importantly, animal and non-animal images were matched *within condition* in their CE and SC values such that animal and non-animal images did not differ from each other in their mean (all t(158) < 0.13, all p > 0.89) or median values (Wilcoxon rank sum test all z < 0.16, all p > 0.87). Images were randomly selected from the larger set of scenes solely based on their image statistics and annotations (animal/non-animal). Animal images contained a wide variety of animals including pets such as dogs and cats but also wildlife, reptiles and fish. Non-animal images consisted of urban, landscape and indoor scenes and contained a variety of objects, ranging from vehicles to household items. Several representative exemplars from each condition are provided in Figure 1*C*.

#### Experimental design

Participants performed the animal vs. non-animal categorization task in a stop signal paradigm (Figure 1*D*). Our motivation for using this paradigm was based on a separate research question inspired by a previous set of studies [52,92], which asked whether the quality of visual input affects decision-making and response inhibition. However, analysis of the behavioral data showed scene clutter to only modulate decision-making on trials with no stop-signal [51–54]. Given that the focus of the current study is on categorization performance rather than response inhibition, trials with a stop-signal presentation are not further discussed. Each participant performed 480 trials in total, divided over 2 separate runs. Each trial lasted 2000 ms and started with a fixation cross of variable duration (500-750 ms jittered with 50 ms intervals), after which a scene (640×480 pixels) was presented for 100 ms. The scene stimuli were back-projected on a 61×36 cm LCD screen that was viewed through a mirror attached to the head coil at ~120 cm viewing distance using Presentation software (Neurobehavioral Systems, Albany, CA, USA). There were two trial types: GO and STOP trials. On GO trials, participants had to indicate whether the stimulus was an animal or non-animal scene before the trial ended (i.e. at maximum within 1250 ms) by pressing one of two buttons. They indicated their response using a hand-held button box with the index or middle finger. If they did not respond in time, a screen displaying the word ‘miss’ appeared for 2000 ms. On STOP trials a beep signal was presented over the headphones indicating that the participant had to withhold their response. At the start of the experiment, the time interval between the stimulus and beep (stop signal delay) was initialized at 250 ms and was adjusted in a staircase procedure based on the stopping performance [93]j. Trials were presented in two randomized sequences that were counterbalanced across participants. Overall, 25% of the scenes (60 animal, 60 non-animal) were shown in STOP trials. The same set of scenes was used in the STOP trials for all participants, and these scenes were excluded from analysis. Thus, all analyses reported here included only the GO trials. For these trials, animal and non-animal scenes were matched in their CE and SC values per condition (means: all t(118) < 1.13, all p > 0.26; medians: all z < 1.10, all p > 0.28).

### fMRI data acquisition

BOLD-MRI data was acquired in a single scanning session, over the course of two runs on a Philips Achieva XT 3T scanner with a 32-channel head-coil located at the University of Amsterdam, The Netherlands. In each run 255 T2*-weighted GE-EPI recordings (TR = 2000 ms, TE = 27.6 ms, FA = 76.1°, SENSE = 2, FOV = 240 mm^2^, matrix size = 80^2^, 37 slices, slice thickness 3 mm, slice gap = 0.3 mm) were made. Breathing rate and heartbeat (using the pulse-oxidization signal recorded from the tip of one of the participant’s fingers) was measured during fMRI acquisition. In addition, a separate functional localizer scan was recorded (317 T2*weighted echo-planar images; TR=1500 ms, TE=27.6, FA=70°, FOV= 240*79.5*240, matrix size= 96^2^, 29 slices, slice thickness 2.5mm, slice gap=0.25) in which participants viewed a series of houses, faces, objects as well as phase-scrambled scenes while pressing a button when an image was directly repeated (12.5% likelihood). These separate localizer scans were used to identify the following regions-of-interest (ROIs): the fusiform face area (FFA), the parahippocampal place area (PPA) and lateral occipital complex (LOC). Finally, a 3D-T1 weighted scan (TE = 3.8 ms, TR = 8.2 ms, FA = 8°, FOV = 256^2^, matrix size = 256^2^, 160 sagittal slices, slice thickness = 1mm) was acquired after the functional runs. This scan was used to register the functional volumes of each run to the structural brain, after which they were registered to standard MNI (Montreal Neurological Institute) space.

### fMRI analysis: animal/non-animal categorization

Analysis was performed using FEAT (FMRI Expert Analysis Tool) Version 6.00, part of FSL (FMRIB’s Software LIbrary, www.fmrib.ox.ac.uk/fsl) and custom Matlab code. The functional data were motion-[94] and slice-time corrected. A temporal median filter was applied to remove low frequencies, after which the data was spatially smoothed with Gaussian kernel at a 5 mm FWHM. The preprocessed scans were subjected to voxel-wise event-related GLM analysis using FILM [95] by convolving the onset times of each trial with a double gamma function to model the hemodynamic response function. We generated explanatory variables (EVs) according to the following conditions: LOW animal, LOW non-animal, MEDIUM animal, MEDIUM non-animal, HIGH animal, and HIGH non-animal. For these EVs, only correct GO trials were included; STOP, omission, and error trials were modeled as separate EVs. In addition, heartbeat and breathing measurements were included in the GLM as nuisance variables. This resulted in an estimate of the BOLD signal for each EV in each voxel, based on which the following contrasts of interest were computed: LOW (animal > non-animal), MED (animal > non-animal) and HIGH (animal > non-animal). As the trial design contained insufficient spacing between trials to estimate a baseline condition, all analyses were conducted on these differential activity measures.

### fMRI analysis: localizer scans

The localizer scans were preprocessed for the purpose of another study conducted within the same experimental session [52]. Again, the data were motion- and slicetime corrected and prewhitened. In addition, they were spatially smoothed using a 4mm Gaussian filter and were temporally filtered by means of a high-pass filter (sigma = 50s). A GLM was fitted with following EVs: for FFA, faces > (houses and objects), for PPA, houses > (faces and objects) and for LOC, intact images > scrambled images. The resulting statistical maps were masked with anatomically-defined regions from the Harvard-Oxford cortical-structural atlas implemented in FSL. For FFA, these were the temporal occipital and occipital fusiform gyrus; for PPA, the parahippocampal gyrus and lingual gyrus, allowing activity to extend posteriorly up to MNI coordinate y=-74; for LOC, the lateral occipital cortex inferior division. Significant voxels within these masks were thresholded at z = 2.3 (FFA and PPA) or z = 3.0 (LOC). For one participant, neither left nor right LOC could be reliably identified: the ROI results for LOC are thus based on 22 instead of 23 participants.

### Statistical analysis: behavioral data

Mean RT and accuracy (percentage correct) was computed for each participant based on the GO trials. Differences in accuracy and RT between the three conditions (LOW, MEDIUM, HIGH) were statistically evaluated using repeated-measures ANOVAs. Significant main effects were followed up by two-tailed, post-hoc pairwise comparisons using a Sidák correction (a variant of Bonferroni correction; Sidak, 1967; Ludbrook, 1991) at α = 0.05. Data were analyzed in Matlab (Mathworks, Natick, MA, USA) and SPSS 22.0 (IBM, Armonk, USA).

### Statistical analysis: whole brain

Whole brain maps were computed by contrasting animal and non-animal responses for each of the three conditions (LOW, MEDIUM, and HIGH). The resulting maps were first pooled across runs (fixed effects) and then across subjects (mixed effects using FLAME1; Woolrich, 2008). after which the following contrasts were run: HIGH (animal > non-animal) > LOW (animal > non-animal); HIGH (animal > non-animal) > MEDIUM (animal > non-animal); and MEDIUM (animal > non-animal) > LOW (animal > non-animal). Results were corrected for multiple comparisons using cluster correction implemented in FSL (z = 2.3, p < 0.05; [99].

### Statistical analysis: ROIs

We additionally examined the contrasts of interest within *a priori* defined regions-of-interest (ROIs) derived from the functional localizer scans (see above) as well as an anatomical mask of V1 derived from the Jülich histological atlas implemented in FSL [100]. Because initial inspection of the results did not indicate major differences between hemispheres, all results are reported for bilateral ROIs. T-values obtained for the contrast of interest were was averaged over voxels within each ROI. The resulting averaged activity values were compared across conditions using one-way repeated-measures ANOVAs. Significant main effects were followed up by post-hoc pairwise comparisons between conditions with a Sidák (REF) multiple comparisons correction at α = 0.05. Data were analyzed in Matlab (Mathworks, Natick, MA, USA) and SPSS 22.0 (IBM, Armonk, USA).

## EEG experiment

### Subjects

Twenty-eight participants (8 males, 19-25 years old, mean = 21.9, SD = 1.9) took part in the EEG experiment. All participants had normal or normal-to-corrected vision, provided written informed consent and received financial compensation. The ethics committee of the University of Amsterdam approved the experiment. Two participants were excluded in preprocessing: one based on behavior (average RT 2 standard deviations from the group average) as well as bad EEG quality (excessive muscle tension reflected in the EEG signal as high frequency noise), and another due to a technical issue which led to duplicate markers being written into the EEG signal.

### Experimental design

Participants viewed the 480 scene stimuli in randomized orders while performing an animal / non-animal speed-accuracy categorization task on the same images used for the fMRI experiment (Figure 1*E*). Stimuli were presented on a 19-inch ASUS monitor with a frame rate of 60 Hz and a screen resolution of 1920 x 1080 pixels. Participants were seated 90 cm from the monitor such that stimuli subtended ~14×10° of visual angle. On each trial, one image was randomly selected and presented in the center of the screen on a grey background for 100 ms. Between trials, a fixation-cross was presented with a semi-randomly chosen duration of either 350, 400, 450, 500 or 550 ms, averaging to 450 ms. Participants searched for animals under either speed or accuracy instructions in randomly alternating blocks that each consisted of 20 trials. Each mini block started with the presentation of an instruction screen displaying either the words ‘QUICK!’ for speeded blocks, or ‘ACCURATE!’ for accuracy blocks for a duration of 5000 ms. In addition, before every trial, the instruction appeared again for 100 ms. Every image was presented twice, once under a speed instruction, and once under an accuracy instruction. After every 120 trials, participants took a short break. Half-way in the experiment, keyboard buttons were switched. Choices and RTs with respect to the start of the presentation of the image were recorded. For both instruction types, participants received feedback on their performance. On the speed trials, participants were presented with “too slow” feedback in case they failed to respond in time (<500 ms), and “on time” when they were quick enough. On the accuracy trials, participants were presented with “correct” and “incorrect” feedback. Stimuli were presented using Presentation software (version 17.0, Neurobehavioral Systems, Inc).

### EEG data acquisition

EEG was recorded with a 64-channel Active Two EEG system (Biosemi Instrumentation, Amsterdam, The Netherlands, www.biosemi.com) at a sample rate of 2048 Hz. The EEG setup was similar to that of our previous studies [36,37,47]. In short, we used caps with an extended 10–20 layout modified with 2 additional occipital electrodes (I1 and I2, which replaced F5 and F6). Eye movements were recorded with additional electrooculograms (EOG). Preprocessing was done in Brain Vision Analyzer 2 (BVA2) and included the following steps: 1) offline referencing to the average of two external electrodes placed on the earlobes; 2) a high-pass filter at 0.1 Hz (12 dB/octave), a low-pass filter at 30 Hz (24 dB/octave), and a notch filter at 50 Hz; 3) automatic removal of deflections larger than 250 mV (after visual inspection, this threshold was raised for some subjects with very high ERP amplitudes); 4) down sampling to 256 Hz; 5) ocular correction using semi-automatic independent component analysis (ICA) followed by visual inspection to identify the components related to eye blinks; 6) segmentation into epochs from -250 to 750 ms from stimulus onset; 7) baseline correction between -200 and 0 ms; 8) automated artifact rejection (maximal voltage 50 μV, minimal/maximal amplitudes - 75/75 μV, lowest activity 0.50 μV); 9), conversion of the obtained ERPs to current source density responses [101]. For the temporal filtering, phase shift-free Butterworth filters implemented in BVA2 were used. Median rejection rate across subjects was 94.5 out of 960 ERP trials (min 8 trials, max 486 trials); in total, 13.6% of the data was rejected. After preprocessing, the ERPs were imported into MatLab (Mathworks, Natick, MA, USA) for statistical analysis.

### Statistical analysis: behavioral data

Choice accuracy and reaction times (RTs) were computed separately for the speed and accurate blocks. Fast guesses (RTs < 150 ms) or RTs > 3 standard deviations from the mean were removed before analysis (mean = 2.2%, SD = 0.9%, min = 0.8%, max = 4.2%). Differences between the LOW, MEDIUM and HIGH condition were tested using two-factor repeated-measures ANOVAs. Significant main effects were followed up by post-hoc pairwise comparisons between conditions or tasks using Sidák multiple comparisons correction at α = 0.05. Data were analyzed in Matlab (Mathworks, Natick, MA, USA) and SPSS 22.0 (IBM, Armonk, USA).

### Drift diffusion modeling

Based on go trial RT distributions of both correct responses and errors, the formal Ratcliff drift diffusion model (DDM) can disentangle the speed of evidence accumulation, ‘drift rate’, the variability of evidence accumulation ƞ, the amount of evidence needed for a decision (a), the starting point of evidence accumulation (z), the variability of this starting point (S_z_). Together these parameters generate a distribution of decision times DT. However, observed reaction times (RT) are also thought to contain non-stimulus specific components such as response preparation and motor execution, which combine in the parameters non-decision time (T_er_), and non-decision time variability (S_t_). In general, DDM assumes that (T_er_) simply shifts the distribution of DT such that: RT = DT+ T_er_ [39,102,103].

To analyze the RT data of the EEG experiment with the drift diffusion model we used a recently developed hierarchical Bayesian estimation of DDM parameters (HDDM), which allows the simultaneous estimation of subject and group parameters and thus requires less data per subject [40]. To gain a deeper insight into how scene complexity affects choice RT, we investigated a model where both the speed of information accumulation (v) and the amount of evidence required to reach a choice (a) were both allowed varying across the three natural scene conditions (LOW, MED, HIGH). Moreover, because previous work has consistently shown that participants require more evidence to reach a choice during accurate instruction trials (as compared to speed trials), evidence requirements (a) was additionally allowed to vary across ‘speed’ or ‘accurate’ instruction trials [104,105]. For this model, three chains of 20,000 samples were generated from the posteriors. In order to assure chain convergence, the first 5,000 samples were discarded (burn), resulting in a trace of 15000 samples for each chain. These chains were then tested for convergence using the Gelman-Rubin statistic, which compares the intra-chain variance of the model to the intra-chain variance of different runs of the same model. All chains were converged and all Rhats were close to 1 [106].

### Statistical analysis: EEG

For each individual subject we computed the average ERP to animal and non-animal scenes in each of the three conditions. Following previous EEG studies on figure-ground segmentation [18,41], ERPs were pooled across a set of occipital and peri-occipital electrodes overlying visual cortex (O1, O2, Oz, POz, PO3, PO4, PO6, PO7, PO8). Per condition, difference waves between animal and non-animal scenes were tested against zero using two-tailed, one-sample t-tests. Difference waves were compared across conditions using the following two-tailed, paired-samples t-tests: HIGH (animal > non-animal) vs. LOW (animal > non-animal); HIGH (animal > non-animal) vs. MEDIUM (animal > non-animal); and MEDIUM (animal > non-animal) vs. LOW (animal > non-animal). Given the large number of statistical comparisons, the results were corrected for multiple comparisons across the two task-instructions, conditions and time-points by means of FDR correction at α = 0.05 using an empirically derived FDR-corrected threshold of *q* = 0.0098).

To compare whether the animal-non-animal ERP amplitude differences in the HIGH condition could be interpreted as the reflection of a slower information processing stream (mediated by feedback), we correlated the speed of information accumulation (drift rate; v) that was estimated for the high complexity scenes with the difference wave amplitude separately for the speed and accuracy task instructions. For this analysis, we selected the interval from first significant deflection of difference wave averaged across speed and accurate instructions (220 ms) until the average peak amplitude (325 ms). We then subdivided this interval into 15 ms bins (7 total) and correlated the average amplitude in that interval with the drift rate (Spearman’s rho, two-sided tests) across participants. Results were corrected for multiple comparisons across time-points using FDR-correction at α = 0.05, yielding *q* = 0.0187 for the accurate trials (no significant time-points were found for the speed trials).

